# Identification of marker-trait associated SNPs for fruit weight and plant height in a wild and domesticated jujube fruit tree (*Ziziphus* spp.) collection using genotyping-by-sequencing

**DOI:** 10.1101/2021.10.30.466592

**Authors:** Nisar Uddin, Harshraj Shinde, Kiflu Tesfamicael, Niaz Ali, Penny J. Tricker, Carlos M. Rodríguez López

## Abstract

*Ziziphus* are economically and nutritionally important fruiting plants that were domesticated in China around 7000 years ago. We identified genetic diversity in 141 accessions of four, different species collected in Pakistan and in China, including wild species of *Ziziphus mauritiana, Z. nummularia* and *Z. oxyphllya*, and both wild and domesticated *Z. jujuba* Mill. Population structure, phylogenetic analysis and principal coordinates analysis based on 10,889 high-quality SNPs derived from genotyping-by-sequencing indicated that the accessions clustered into two major groups. The wild Pakistani *Z. jujuba* and *Z. nummularia* exhibited higher genetic diversity and polymorphic information content (0.37 and 0.23 respectively) than other species. We further conducted a genome-wide association study and discovered six highly significant marker-trait associations for fruit weight and plant height in this population. Our study provides important information for future breeding of *Ziziphus* species.

## 1. Introduction

*Ziziphus* is a genus of around 200 species of spiny shrubs, small trees, and vines in the buckthorn family, Rhamnaceae. Species are distributed throughout the world in warm-temperate and subtropical regions. *Zizphus* are grown for their jujube fruits in ∼50 countries, including in arid and semi-arid regions of Australia, India, Iran, Israel and Pakistan, but more than 90 % of all production is currently concentrated in China. Although breeding for *Ziziphus* cultivars via selection has been practiced for more than a thousand years, the process of cross-breeding is difficult due to their tiny flowers that are hard to emasculate (Wang et al. 2014), and trees are mostly propagated by grafted cuttings (Xu et al. 2016).

The fruits of *Ziziphus* plants are highly nutritious (Abdoul-Azize 2016) and they are sources of carbohydrates, minerals (iron and potassium), phenolic compounds and vitamins (Rashwan et al. 2020). They also present medicinal properties and are widely used in local and traditional medicines (Ahmad et al., 2013; 2016). Pharmacological activities of different Ziziphus species have been identified, such as the anti-ulcerogenic activity of *Z. lotus*, anti-cancer proliferation properties of *Z. jujuba* and antimicrobial activity of *Z. mauritiana*. With these characteristics, *Ziziphus* fruits can be considered ‘super foods’ with potential for expanded production and consumption (Liu et al. 2020). Globally, *Ziziphus* are becoming increasingly popular to grow because of their ability to adapt to a broad range of climatic conditions (Jiang et al. 2020). They yield well in arid and semi-arid regions, where other fruit trees fail to grow, and are relatively tolerant of drought, salinity and high temperature stress (Liu et al. 2020).

To explore and capture the genetic diversity available for selection and breeding for improvement of quantitative traits in such species, one efficient method is genome-wide association study (GWAS) where polymorphic markers from a diverse collection, rather than an inbred population, are associated with phenotypic traits. Such studies also allow the exploration of markers-trait associations using the wild relatives of domesticated cultivars that may still harbor beneficial alleles lost during domestication and the narrowing of a gene pool (McCouch 2013), and panels of accessions from different geographic origins where plants may have adapted to different climatic conditions. In order to test the available genetic variation in a collection and perform GWAS, a high, genome-wide, marker density is required. Among different molecular makers, single nucleotide polymorphisms (SNPs) are widely used and important molecular markers (Mammadov et al. 2012). In plants, genotyping by sequencing (GBS) using high-throughput sequencing has proved to be an effective method for the development and genotyping of a large number of SNP markers (Peterson et al. 2014; Torkamaneh, et al. 2016; Belzile, 2016; Pereira-Dias et al. 2019; He et al. 2014).). GBS has been successfully used in many crop species including maize, barley, wheat, rice (Poland and Rife, 2012) and, recently, in *Z. jujuba* Mill. for molecular markers discovery in a core collection (Chen et al. 2017) and for GWAS of domestication and fruit traits (Gou et al., 2020; Guo et al., 2021; Hou et al. 2020). Molecular markers for any identified quantitative trait loci (QTL) can then be used for marker-assisted selection in plant breeding programs (He et al. 2014) to incorporate alleles from diverse, genetic resources.

In our current research, we report the successful application of genotyping by sequencing in a 141accession panel of five different *Ziziphus* species and origin groups, generating more than 10,000 high quality SNP markers. These were used to investigate the population structure, genetic diversity and phylogeny of the collection. We also performed GWAS to associate markers with phenotypic and agronomic traits. Our results provide a valuable foundation for the breeding and genetic improvement of *Ziziphus* crop species.

## 2. Materials and Methods

### 2.1. Samples collection

In total, 189 leaf samples were collected from mature *Ziziphus* trees in different agro-ecological regions of Pakistan and China during two consecutive years, 2017-2018 (Supplemental Table S1). Samples of the species, *Ziziphus nummularia* (84 samples), *Z. jujuba* var. spinosa (87 samples) and *Z. oxyphllya* (5 samples) were collected from Pakistan, whereas *Z. mauritiana* (5 samples) and the cultivar *Z. jujuba* var. jujuba (8 samples) were collected from China.

### 2.2. Phenotypic trait measurements

11 quantitative traits were measured at the time of collection. Number of branches (BR) were visually counted. Plant height (PH), leaf length (LL), leaf width (LW), leaf thickness (LT), petiole length (PL), internode length (InL) and stem diameter (SD) were estimated using a measuring tape. Fruit weight (FtW) was calculated as the average weight of 10 mature fruits from the 4^th^ and 5th branches. Maximum fruit diameter (FtD) and fruit length (FtL) of these fruits were measured using vernier calipers. Phenotypic data are summarized in Supplemental Table S2.

### 2.3. Nucleic acid extraction, GBS library preparation and SNP analysis

Leaf samples collected for each accession were rapidly frozen in liquid nitrogen and stored at - 80°C until used. Samples were powdered using pestle and mortar in liquid nitrogen and genomic DNA was extracted using the modified CTAB (cetyl trimethyl ammonium bromide) method (Allen et al. 2006). The quality and concentration of the extracted genomic DNA was assessed by spectrophotometer (Thermo Fisher scientific, USA) and 1% agarose gel electrophoresis.

GBS libraries were prepared as described by Kitimu et al (2015) using DNA extracted for all 189 samples. Briefly, 100 ng of genomic DNAs were enzymatically digested using *EcoR*I and *Msp*I (NEB, Ipswich MA, U.S.A) and ligated to *EcoR1* and *Msp1* adaptors. Each sample was assigned a unique barcode and PCR amplicons of all samples were equimolarly pooled together into two libraries, each library containing 96 samples (plus 3 water controls). PCR amplicons were cleaned using AMPure XP bead with equal ratios. The quality of amplicons was measured using a 5200-fragment analyzer (Agilent, USA). Libraries were sequenced using the Illumina Hi-Seq 2500 platform with output of 150 bp single end reads. Single nucleotide polymorphism (SNP) calling was performed using the reference-based GBS pipeline (Torkamaneh, Laroche, and Belzile 2016) as described by Tesfamicael et al. (2020). Before SNP calling, sequence reads were filtered for those harboring identical matches to barcodes, minimum kmer count (< 10) and kmer Length (< 20). To deduce the genomic position of read tags, sequence reads were mapped to the reference genome NAFU_Zjuj_v2.0 (*Ziziphus jujuba* Junzao, GCA_001835785.2).

### 2.4. Population structure, genetic and phylogenic analyses

Population structure was evaluated with 10,899 SNP markers via STRUCTURE 2.1 software (Pritchard 2007). The number of hypothetical subpopulations (*K*) was set from 2-5. The best *K*-value was determined using STRUCTURE HARVESTER (Earl and Vonholdt, 2012) based on delta *K*(Δ*K*) and maximum log likelihood L(*K*). For this analysis, the length of the burn-in period and number of the MCMC replications after burn-in were set to 50,000 iterations and 50,000, respectively. Major allele frequency, genetic diversity, heterozygosity and polymorphic information contents were calculated using the Power Marker Software (Liu and Muse 2005) with default settings. Phylogenetic tree and principal component analysis (PCA) were analyzed using TASSEL 5 (Bradbury et al. 2007).

### 2.5. Genome-wide association study (GWAS)

A mixed linear model principal components analysis incorporating population structure (Q) and kinship (K) was built using TASSEL 5 (Bradbury et al. 2007) and quantile-quantile plots were generated in the R environment using gplots and examined for the expected and observed distribution of each trait (Supplemental Fig. S1). Marker-trait associations were predicted for 11 traits using the Genome Association and Prediction Integrated Tool (GAPIT) in the R environment and for the generation of Manhattan plots (Lipka et al. 2012). For marker-trait associated SNPs (MTAs) analysis the significance threshold was adjusted to p= 0.05 after applying a false discovery rate correction (Benjamini and Hochberg 1995). MTAs below the threshold p= 0.05 were considered significant. Sequences of significant MTAs were BLAST in the *Ziziphus jujuba* cv. Dongzao reference genome (ZizJuj_1.1 RefSeq GCF_000826755.1). Associated genes annotated in NCBI Release 101 located within the 39 kbp average marker distance in this genome were identified.

## 3. Results

### 3.1. SNPs identification and analysis

GBS generated a total of 471,854,625 150 bp length reads. Our reference genome-based SNP calling initially generated 509,250 SNPs and 38.6 % of reads were mapped to the reference genome of *Ziziphus jujuba* cv. Junzao. 48 samples were eliminated because of the high numbers of missing SNPs, leaving 141 samples for further analysis. After filtering for SNPs with missing values, 10,889 high-quality SNPs remained. Of these, 10,167 (93%) physically mapped to one of the 12 chromosomes of the reference genome and the remaining 722 (7%) were mapped to scaffolds. The average number of SNPs per chromosome was 848 with the highest number (1429) on chromosome 11 and the lowest (541) on chromosome 8. C/T transitions were the highest frequency SNPs (15.09 %), followed by A/G transitions (14.29 %) and G/A transversions (12.43 %) (Supplemental Table S3).

### 3.2. Phylogenetic and principal component analysis

Phylogenetic analysis (Figure 1a) separated the studied samples into two main clades. One composed by *Z. jujuba* accessions from Pakistan and China, with Chinese jujube forming a distinct sub-clade within the *Z. jujuba* group. The second clade contained accessions from *Z. nummularia, Z. mauritiana*, and *Z. oxyphylla*, with the former two, not forming clear subclades within the *Z. nummularia* accessions. PCA analysis showed that the first three principal components explained 69% of the total variability captured by the 10,889 high-quality SNPs used in this analysis (i.e, PC1 = 52%; PC2 = 10%; PC3 = 7%). In this case, accessions from Z. *jujuba sensu lato* formed a dispersed cluster that occupied the right two quadrants of the Eigen space, *Z. nummularia*, and *Z. oxyphylla* formed a more compact cluster occupying the bottom left quadrant, while *Z. mauritiana* formed a separate cluster between the two main clusters (Figure 1b).

**Figure 1.**
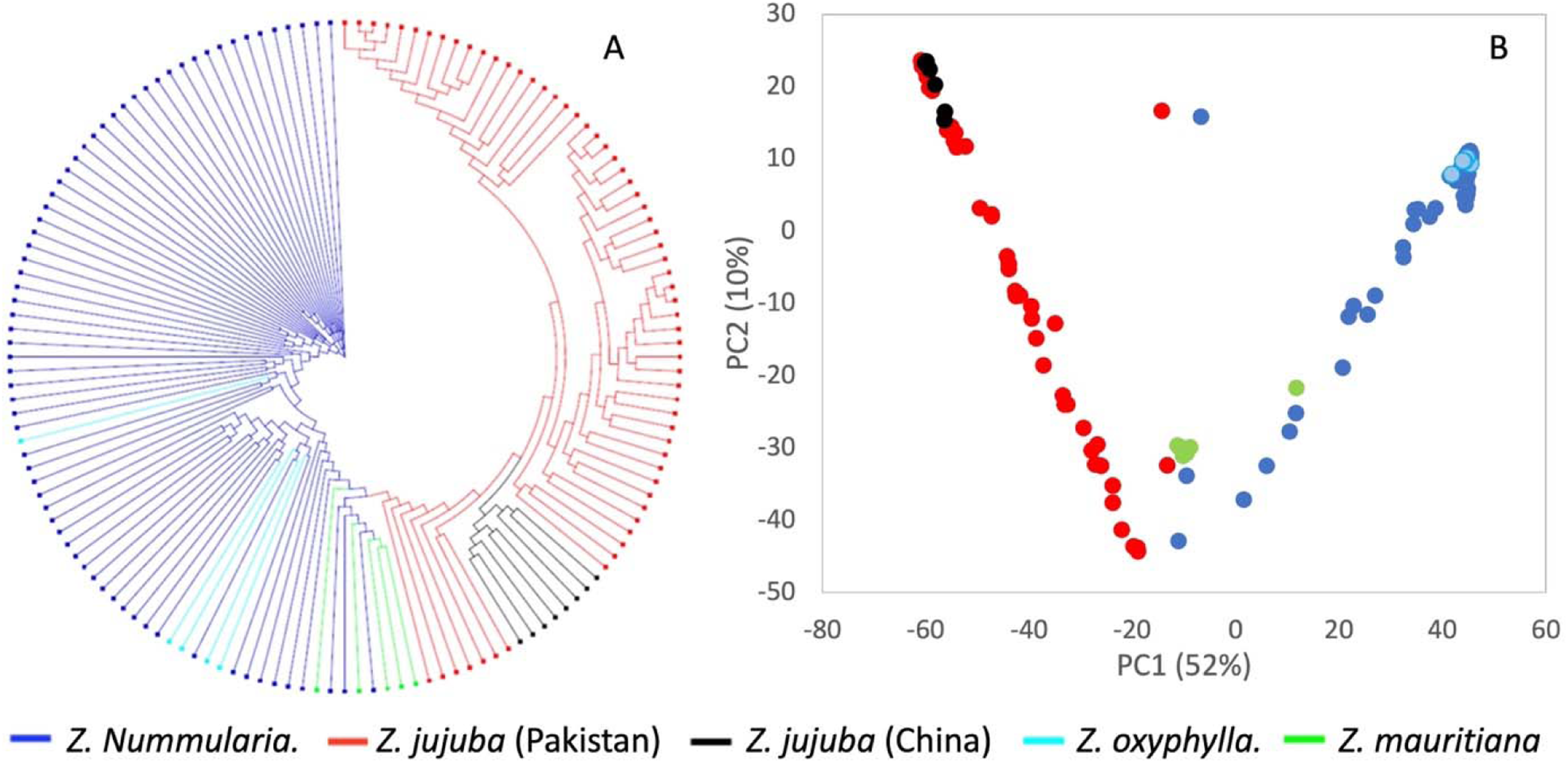
Phylogenetic tree and Principal Co-ordinates Analysis of the *Ziziphus* collection comprising 141 genotypes. Both analyses were carried out using 10,889 high-quality SNPs generated using genotyping by sequencing.

### 3.3. Population structure and phenotypes within clusters

Population structure analysis for GWAS exhibited the highest ΔK at K = 2, where the five *Ziziphus* species and origin groups divided into two clusters, with some admixture among groups (Figure 2). Cluster 1 included all genotypes of *Z. nummularia* and *Z. oxyphylla*, whereas cluster 2 comprised all the *Z. mauritiana* samples and all members of *Z. jujuba* from both China and Pakistan. Plant height was strongly differentiated by cluster with the average height of trees in cluster 2 much taller than in cluster 1, but the range of heights also much more spread reflecting species’ differences within cluster 2 (Figure 3). Other architectural traits (branching, stem diameter) were generally greater in cluster 2, again with a larger range. Leaves were also longer in cluster 2 genotypes, although thicknesses and widths were similar in both clusters. The range of fruit diameters and lengths was bigger in cluster 2, although the median fruit weight was higher in cluster 1.

**Figure 2.**
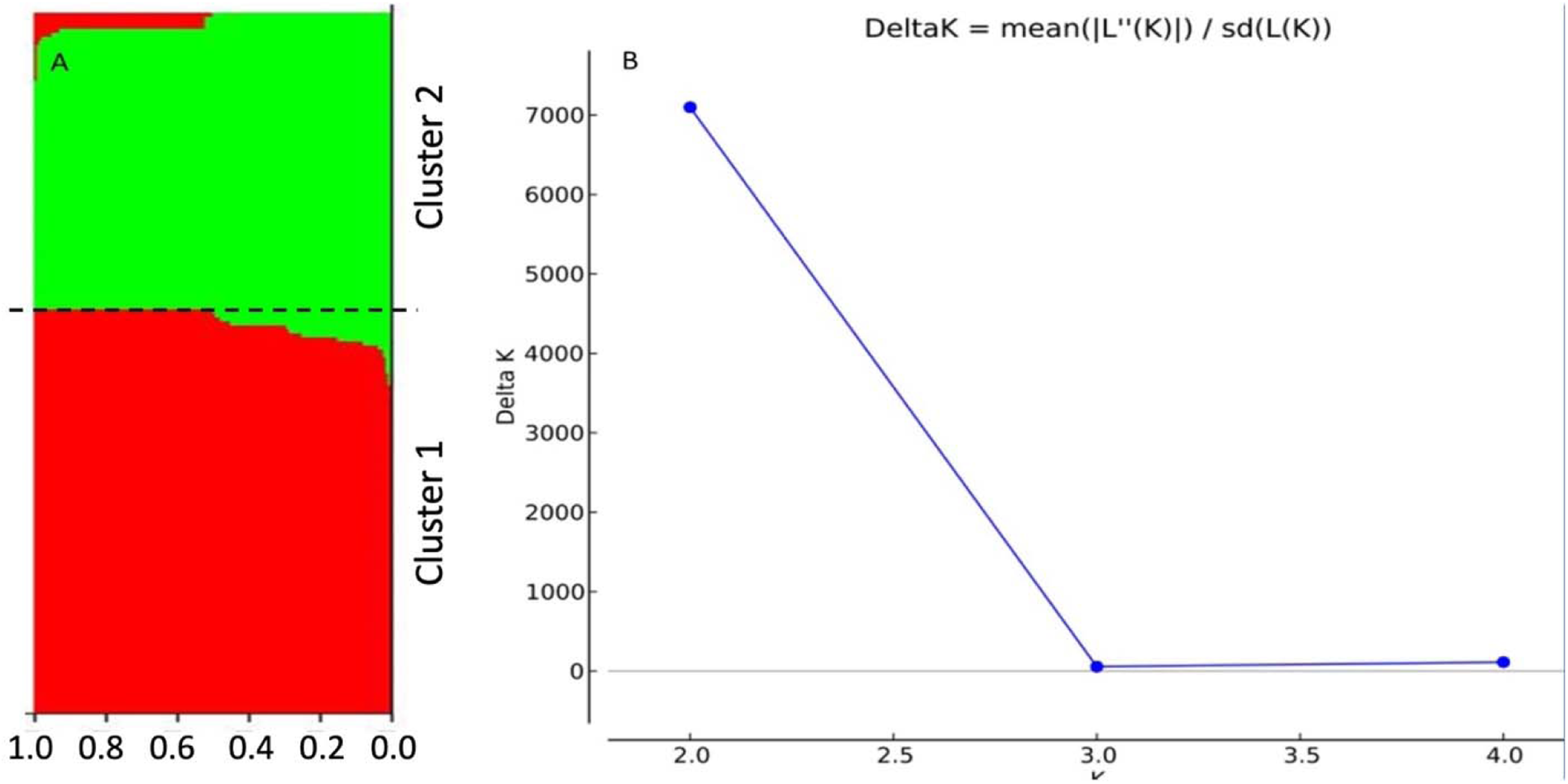
Population structure analysis of the collection. (A) Population structure based on *K* = 2. Each colored bar is an accession, and the width of the bar is an estimate of the proportion of its membership in each cluster. Cluster 1 includes all accessions from *Z. nummularia* and *Z. oxyphylla*. Cluster 2 includes all accessions from *Z. mauritiana Z. jujuba* both from China and Pakistan. B) Evanno plot of Δ*K* calculated from *K* ranging from 2 to 5.

**Figure 3.**
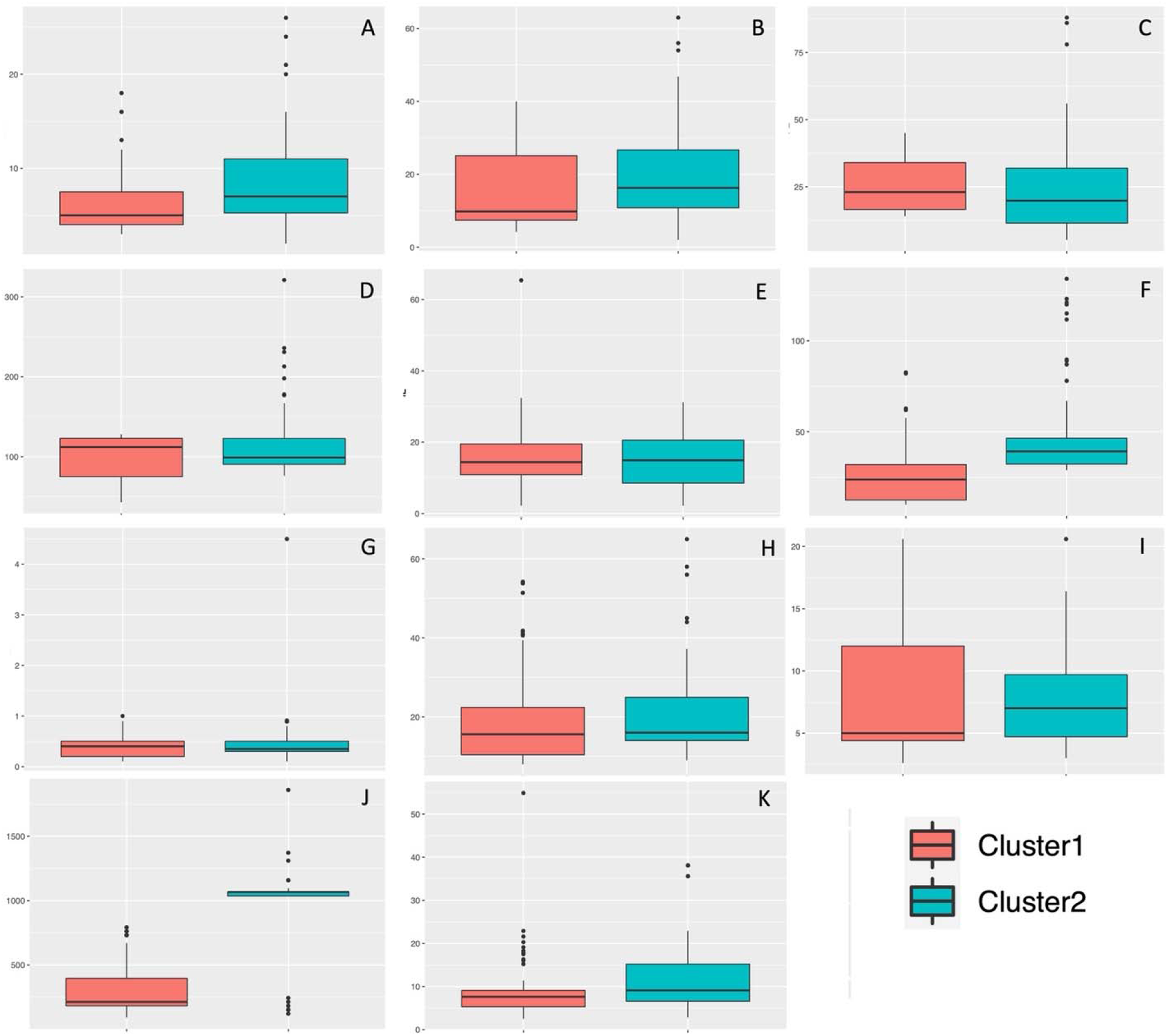
Boxplots showing medians and range of phenotypic traits within each of the two population clusters for: Branching (A), Fruit diameter (B), Fruit length (C), Fruit Weight (D), Internode length (E), Leaf length (F), Leaf thickness (G), Leaf width (H), Petiole length (I), Plant height (J) and Stem diameter (K).

### 3.4. Genetic diversity of the *Ziziphus* species

The wild species collected in Pakistan, *Z. nummularia* and *Z. oxyphylla*, exhibited higher heterozygosity (He) than either wild or domesticated *Z. jujuba* or *Z. mauritiana* (Table 1). The major allele frequency (MAF) was high in all species, ranging from 0.64 in Pakistani *Z. jujuba* to 0.94 in Chinese *Z. jujuba*, but the polymorphic information content (PIC) was variable between the species and origins, highest (0.37) for Pakistani *Z. jujuba* and low (0.07) for *Z. oxyphylla* and for Chinese *Z. jujuba* (0.08).

**Table 1.**
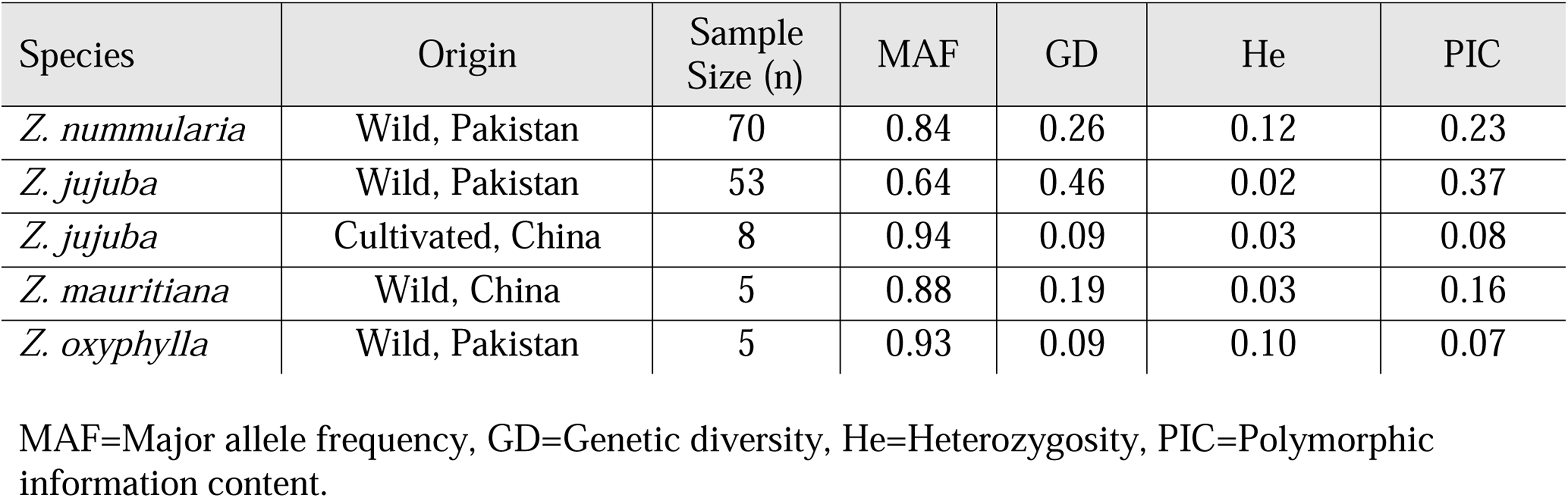
Genetic diversity analysis of different *Ziziphus* species and origin groupings, based on ∼10,900 SNP markers.

| Species | Origin | Sample Size (n) | MAF | GD | He | PIC |
| --- | --- | --- | --- | --- | --- | --- |
| *Z. nummularia* | Wild, Pakistan | 70 | 0.84 | 0.26 | 0.12 | 0.23 |
| *Z. jujuba* | Wild, Pakistan | 53 | 0.64 | 0.46 | 0.02 | 0.37 |
| *Z. jujuba* | Cultivated, China | 8 | 0.94 | 0.09 | 0.03 | 0.08 |
| *Z. mauritiana* | Wild, China | 5 | 0.88 | 0.19 | 0.03 | 0.16 |
| *Z. oxyphylla* | Wild, Pakistan | 5 | 0.93 | 0.09 | 0.10 | 0.07 |

MAF=Major allele frequency, GD=Genetic diversity, He=Heterozygosity, PIC=Polymorphic information content.

### 3.5. Genome-Wide Association Study (GWAS)

The GWAS was performed for 11 different phenotypic traits and, using a conservative approach, six significant marker-trait associations (MTAs) were discovered for two of these traits (Figure 4). Four significant MTAs were found for fruit weight on three chromosomes, chr. 3, chr. 8 and chr. 10. For plant height two significant MTAs were identified, one on chr. 3 and one on chr. 8. The associations at the marker 8_1006069 were significant for both traits with a positive allelic effect of 9.09 % on plant height and negative allelic effect of -38.22 % on fruit weight, indicating the importance of this marker and an inverse correlation between these two traits at this locus.

**Figure 4.**
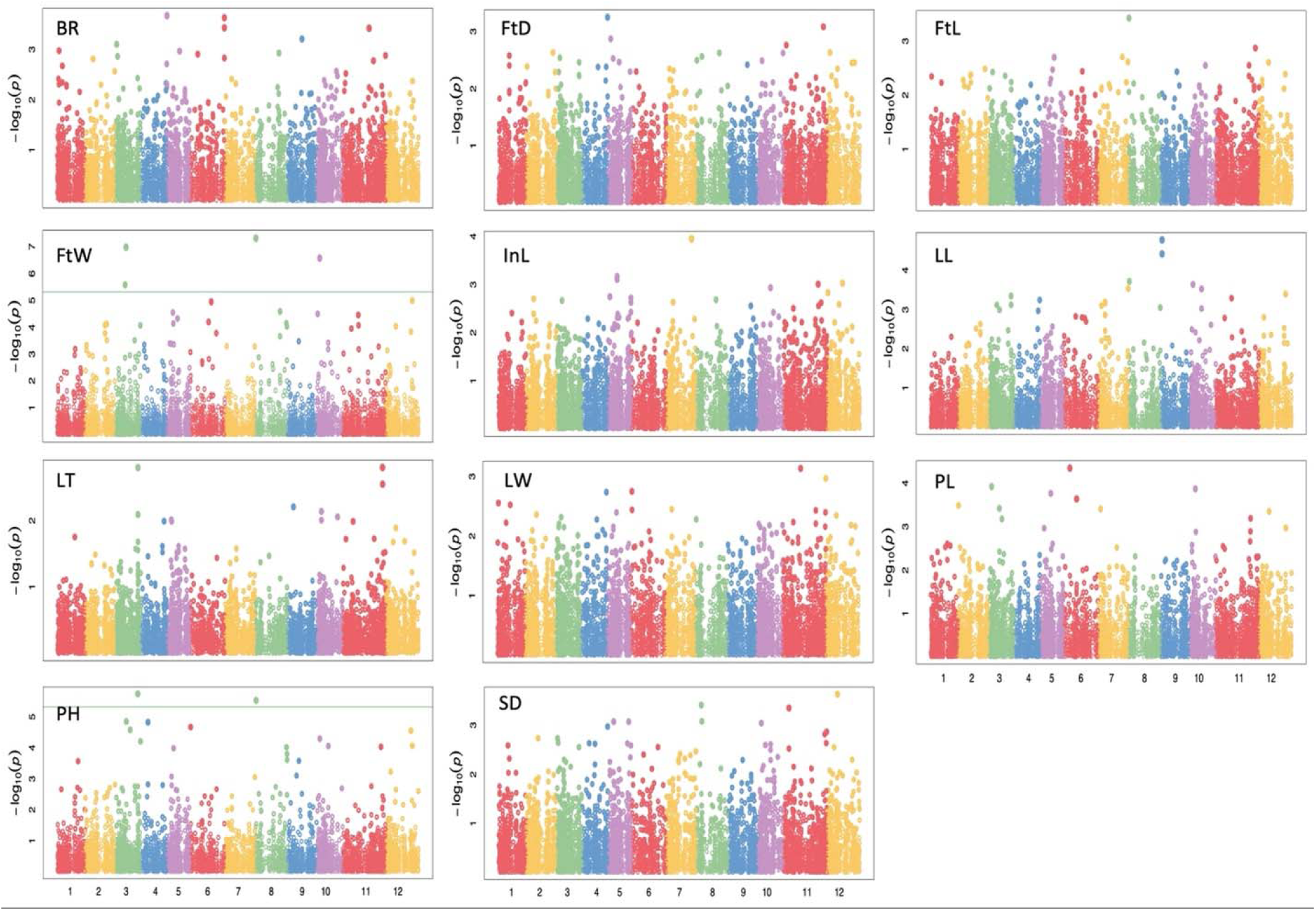
Manhattan plots of marker-trait associations for Branching (BR), Fruit diameter (FtD), Fruit length (FtL), Fruit Width (FtW), Internode length (InL), Leaf length (LL), Leaf thickness (LT), Leaf width (LW), Petiole length (PL), Plant height (PH) and Stem diameter (StD) traits. Green lines denote the genome-wide significance threshold.

No genes are yet annotated in the annotated v1.1 reference sequence for *Z. jujuba* at this QTL (annotated genes at loci are listed in Supplemental Table S4) and only one of all the significant markers (3_10271794 for fruit weight) was a direct hit in an annotated gene (LOC107413626) in this reference. The remaining three significant MTAs for fruit weight each had high and positive effects of between 41-43.5 %. Marker 3_20731903 for plant height had a high negative effect (−20.91 %) and a much lower major allele frequency (0.025) in this collection.

## 4. Discussion

For this investigation, we successfully applied GBS technology to generate high-quality SNPs across four *Ziziphus* species. The number of SNPs generated in our study (10,889) is higher than in many similar studies, including the previously reported 4,680 SNPs from GBS in a 150 member *Z. jujuba* core collection (Chen et al. 2017) and the 4,651 SNPs found in the extended, 180-accession *Z. jujuba* panel (Hou et al. 2020). This suggests, not only that GBS and reference sequence-based alignment is an efficient approach for SNPs identification in related *Ziziphus* species, but also that the addition of accessions from different species and origins allowed us to capture previously untapped molecular diversity that could prove valuable for marker-assisted approaches to both conservation and breeding.

Both the phylogenetic and population structure analysis of our collection grouped the accessions into two major clades; one clade comprised all *Z. jujuba* accessions and the other included all *Z. nummularia* and *Z. Oxyphylla*. Although *Z. mauritiana* accessions clustered with *Z. jujuba* in population structure analysis, phylogenetic and PCA analysis were more sensitive, revealing that these accessions were distinct in comparison with either the *Z. jujuba* or the *Z. oxyphylla*, with some admixture with *Z. nummularia*. These results agree with the findings of Uddin et al. (2021) for a larger collection of 200 genotypes (including the 141 accessions studied here) characterized using three highly polymorphic SSRs. The country of origin of the *Z. jujuba* did not influence the population structure and, although the limited number of Chinese *Z. jujuba* cultivars in this collection clustered together, this clade was within the wider *Z. jujuba* clade, suggesting that the molecular diversity in the wild *Z. jujuba* from Pakistan could be a useful source of diverse alleles for the cultivated varieties. Although the clades broadly separated along species lines, this was not complete, with markers sufficiently shuffled to complete the association study using a mixed linear model, with a conservative correction, and low false discovery rates for the MTAs. Interestingly, the *Z. nummularia* and *Z. oxyphylla* clade was the most mixed between the species, but with the furthest separation from *Z. jujuba*, suggesting that they may have a unique ancestry.

The highest genetic diversity and polymorphic information content in our collection was measured in the wild *Z. jujuba* from Pakistan, whereas the lowest diversity was found in the cultivated and domesticated *Z. jujuba* from China. Fresh jujube (*Z. jujuba* var. jujuba) was domesticated from wild, dry jujube (*Z. jujuba* var. spinosa) in China, and the wild relative progenitor of sweet *Z. jujuba* is *Z. acidojujuba*. Huang et al. (2016) resequenced 31 accessions of *Z. jujuba* var. spinosa and Guo et al. (2021) resequenced 493 *Z. jujuba* and *Z. acidojujuba* accessions and 350 wild (var. spinosa), semi-wild (var. spinosa) and cultivated (var. jujuba) *Z. jujuba* accessions, all collected in China. Comparing whole genome assemblies (Huang et al., 2016) and using GWAS (Guo et al., 2020; 2021) the chromosomal regions of selective sweeps and important traits and genes for domestication were identified. Despite large differences in genome size between fresh and dry jujube (Huang et al., 2016), relatively little whole genome nucleotide diversity between the wild progenitor (*Z. acidojujuba*), wild, semi-wild (var. spinosa) and domesticated, cultivated *Z. jujuba* (Guo et al., 2020; 2021) was found in these studies. Our finding that relatively high genetic diversity has been retained in the *Z. jujuba* accessions from Pakistan in comparison with those from China is, therefore, surprising, but is consistent with previous findings for this collection (Uddin et al., 2021). This result might have reflected the smaller number of *Z. jujuba* from China in the panel, but polymorphic information content was also higher in the small number of *Z. mauritiana* than in the Chinese *Z. jujuba*. Alternatively, it could indicate that *Z. jujuba* has been under lower selection pressure (either natural or artificial) in Pakistan than in China. Most significant MTAs identified here did not coincide with previously reported regions of selective sweeps or validated candidate genes for domestication that included those for fruit shape and weight (Guo et al., 2020; 2021). The marker 8_1006069 that was significantly associated with both fruit weight and plant height in our GWAS is in the region identified by Guo et al. (2020) for bearing-shoot length and number of leaves, suggesting that selection for plant architecture during domestication may underly this QTL.

Although the large number of high quality SNPs and population structure of this panel allowed us to confidently identify significant MTAs through GWAS, differences in genome size (Huang et al., 2016) and linkage disequilibrium decay (Guo et al. 2020; 2021) between different *Ziziphus* species have been reported and, using a reference-based alignment approach, we were unable to assess potential structural variations between species’ genomes. Thus, although reference-based alignment was a practical option, our study is limited by this approach in comparison with full, pan-genome approaches. Although we have listed genes annotated and positioned within the region of the QTL in the reference genome (Supplemental Table S4), we have no knowledge of whether those positions vary, or whether these genes are present at those positions in the other *Ziziphus* genomes included here. Although the markers identified can be used for marker-assisted selection, further work would be required to validate any candidate genes. Nonetheless, to the best of our knowledge, the MTAs identified in this study were novel (although some similar fruit traits were studied using GWAS by Hou et al. (2020) in *Z. jujuba*) and they had high allelic effects. Utilizing the different species and origins resulted in a wide range of trait phenotypes for the majority of traits that allowed us to explore novel genetic associations. For most traits this range was greater in cluster 2 that included all the *Z. jujuba* and *Z. mauritiana* accessions. This, again, indicated, that genetic diversity has been retained in *Z. jujuba* in this collection, but also the wider range of phenotypes allowed us to uncover novel MTAs for important traits. For example, all the *Z. jujuba* of Chinese origin were relatively short, all the *Z. mauritiana* relatively tall, with the remainder of accessions from other species and origins distributed between. Likewise, the highest fruit weights were exclusively observed in the Chinese *Z. jujuba*, the lowest in *Z. oxyphylla*, with the remainder between. The evident genetic determination underlying these traits allowed us to identify MTAs that would likely have been masked in a single species or origin collection.

Our results demonstrate that the genetic diversity among different *Ziziphus* species can be of value for ongoing crop improvement programs and suggest that the continued conservation and development of additional genetic and genomic resources for the orphan crops in this genus will be worthwhile.

## Author Contributions

CMRL and NU conceived the study. CMRL and NA acquired funding and administered the project. NU performed the phenotypic characterisation. NU, KT, HS and PT performed molecular biology and bioinformatics analysis. HS, NU and PT wrote the manuscript. CMRL critically revised the manuscript.

## Acknowledgments

The authors thank multiple members of the EEGG lab for valuable suggestions to improve this manuscript.

## Supplementary Material

**Supplementary table S1**. Collection site and population structure cluster of the 189 *Ziziphus* accessions studied. (NZNL-N-*Ziziphus Ziziphus nummularia*, NZJL-J-*Ziziphus jujuba* (Pakistan), ZOXL-OX-*Ziziphus oxyphylla*, ZML-M-*Ziziphus muaritania* and *CHZL-C-Ziziphus jujuba* (China)). Accessions with high number of missing SNPs were not included in the analysis and therefore not assigned to a cluster.

**Supplemental Table S2**. Phenotypic data for the 189 Ziziphus accessions included in this study. Species and origins: N = *Ziziphus nummularia*, O = *Z. oxyphylla*, M = *Z. mauritiana*, J = *Z. jujuba* (from Pakistan), C = *Z. jujuba* (from China).

**Supplemental Table S3**. List of 10,889 filtered, high quality single nucleotide polymorphisms discovered in 141 genotypes.

**Supplementary Table S4**. Marker-trait associations. SNP markers associated with the traits identified using GWAS. Associated genes were called when present within the calculated average marker distance in this genome (39 Kbp). Bold font denotes a SNP within an annotated gene.

**Supplemental Figure S1**. Quantile-quantile scatterplots from the mixed linear model analysis for each of the phenotypic traits measured (Branching (BR), Fruit diameter (FtD), Fruit length (FtL), Fruit Width (FtW), Internode length (InL), Leaf length (LL), Leaf thickness (LT), Leaf width (LW), Petiole length (PL), Plant height (PH) and Stem diameter (SD)).

### Data Availability Statement

The raw sequencing data are available in the NCBI-SRA database under the bioproject accession number PRJNA679591.

